# Nicotinamide Riboside Supplementation Ameliorates Mitochondrial Dysfunction and Neuronal Loss in POLG Mutant Midbrain Organoids

**DOI:** 10.1101/2023.11.08.566203

**Authors:** Tsering Yangzom, Anbin Chen, Gareth John Sullivan, Kristina Xiao Liang

## Abstract

Mitochondrial dysfunction is associated with many neurodegenerative disorders and is particularly prominent in conditions tied to *POLG* mutations. *POLG* encodes DNA polymerase gamma vital for mitochondrial DNA replication. Employing 3D human pluripotent stem cell-derived midbrain organoids (hMOs), harbouring *POLG* mutations, this study explores their differentiation, transcriptional alterations, and underlying pathways of neurodegeneration associated with *POLG* mutations. The generated hMOs displayed midbrain specificity and, at three months, a reduced diameter, suggesting growth challenges from *POLG* mutations. A reduced presence of dopaminergic neurons, particularly in DA2 and ventral midbrain classes, was evident. Intriguingly, post-treatment with 1 mM Nicotinamide Riboside (NR), an NAD^+^ precursor, the organoids demonstrated an increased count of DA and VMN neurons and an elevated gene expression, especially in processes crucial to mitochondrial and synaptic functions. Our findings spotlight NAD^+^ supplementation has potential therapeutic value in addressing POLG-associated neuronal and mitochondrial deficits. Moreover, the unique insights garnered from single-cell RNA sequencing, and enrichment analyses further emphasize the significance of mitochondrial disturbances and potential interventions for POLG-related neurodegenerative conditions. In summary, we underscore the transformative potential of NAD^+^ in managing neurodegenerative diseases associated with *POLG* mutations. It also establishes the utility of *POLG* mutant hMOs as a potent research model.

## Introduction

Mitochondria, often referred to as the cellular “powerhouses”, play a pivotal role in cellular bioenergetics, especially in neurons where energy demands are exceptionally high. Dysfunctional mitochondria are implicated in numerous neurodegenerative disorders, posing a significant threat to the health, and functioning of neural circuits. A major contributing factor to this is the presence of mutations in the POLG gene, which encodes DNA polymerase gamma, an enzyme essential for mitochondrial DNA replication. These mutations not only destabilize mitochondrial DNA (mtDNA) but also give rise to neuronal anomalies, the hallmark of many neurodegenerative conditions.

The complexity of the human brain and cellular processes presents a formidable challenge to researchers. Notable advancements have been achieved using human induced pluripotent stem cells (iPSCs) to generate region-specific neuronal subtypes (1, 2). These advances have yielded invaluable insights into human neural development and various neurodevelopmental disorders (2, 3, 4, 5). However, conventional two-dimensional cell cultures often fall short of replicating the true complexity of the brain (6, 7, 8). Recognizing these limitations, brain organoid technology has emerged as a revolutionary approach. It leverages the self-organizing capabilities of iPSCs, closely mimicking key milestones in early brain development (9, 10, 11, 12, 13). Emphasizing the significance of this shift towards three-dimensional (3D) modelling, several distinct organoid models have been developed, each targeting specific aspects of brain functionality and spatial organization. These models have been instrumental in advancing our understanding of disorders including Parkinson’s disease (PD), and microcephaly (14, 15, 16, 17, 18, 19).

Amidst the challenges presented by mitochondrial dysfunction linked to *POLG* mutations, the search for effective therapeutic solutions continues. One intriguing solution that has garnered considerable interest is nicotinamide riboside (NR), a precursor to the coenzyme NAD^+^. Given NAD^+^’s critical role in cellular metabolism, its potential neuroprotective properties, especially in the context of compromised mitochondrial health, have gained prominence.

This investigation shows the impact of *POLG* mutations within iPSC-derived midbrain organoids and explores the therapeutic potential of NAD^+^ supplementation using NR. This study’s overarching goal is to shed light on the intricacies of POLG-associated pathology and potentially pave the way for effective interventions in the management of neurodegenerative disease.

## Methods

### Ethics Approval

This study received approval from the Western Norway Committee for Ethics in Health Research (Approval Number: REK nr. 2012/919).

### iPSCs Culture

Patient-derived iPSCs, generated from parental fibroblasts with *POLG* mutations, notably one homozygous for c.2243G>C; p.W748S (WS5A), were cultured following methods previously established (20, 21, 22, 23). Disease-free control iPSCs were reprogrammed using fibroblast lines Detroit 551 and CCD-1079Sk (24, 25, 26). For hMO differentiation, we also incorporated human embryonic stem cell line 360 from the Karolinska Institute, Sweden (27) as a reference control.

The iPSCs and ESCs were maintained on 6-well plates coated with Geltrex (#A1413302, Thermo Fisher Scientific). For the initial 24 hours following seeding, they were cultured in Essential 8 Basal medium (#A1516901, Thermo Fisher Scientific) enriched with Y-27632 dihydrochloride Rock Inhibitor (#1254, Biotechne Tocris). Daily medium changes were performed, and upon reaching a confluency of approximately 70%-80%, iPSCs were passaged using 0.5mM EDTA (#15575038, Thermo Fisher Scientific). For quality control, we routinely checked cell cultures for the presence of mycoplasma using the MycoAlert™ Mycoplasma Detection Kit (#LT07-218, Lonza).

### hMOs Generation and Cultivation

iPSCs and ESCs were propagated on 6-well plates coated with Geltrex (#A1413302, Thermo Fisher Scientific). These cultures were grown in Essential 8 Basal medium (#A1516901, Thermo Fisher Scientific), which was augmented with Y-27632 dihydrochloride Rock Inhibitor (#1254, Biotechne Tocris) during the initial 24 hours following seeding. We ensured that the culture medium was replaced daily. Once the iPSCs reached a confluency of approximately 70%-80%, they were dissociated using 0.5mM EDTA (#15575038, Thermo Fisher Scientific). As part of our routine quality control process, the presence of mycoplasma in the cultures was regularly screened using the MycoAlert™ Mycoplasma Detection Kit (#LT07-218, Lonza).

### Generation and Maintenance of hMOs

The generation and maintenance of hMOs were followed as described previously (28). Briefly, iPSCs (~70% confluence) were plated on Geltrex-coated dishes. The next day, we used a neural induction solution enhanced with SB431542 (#1614, Tocris Bioscience), NAC (#A7250, Sigma-Aldrich), and Compound C (#US1171261 1MG, EMD Millipore). This combined with a CDM containing IMDM (#21980 065, Thermo Fisher), F12 Nutrient Mixture with GlutaMAX™, BSA Fraction V (#EQBAC62 1000), Lipid 100 X (#11905 031, Thermo Fisher), 1-thioglycerol (#M6145 25ML, Sigma-Aldrich), insulin (#11376497001, Roche), and transferrin (#10652202001, Roche). By day 5, after reaching the neural epithelial phase, cells were moved using collagenase IV to untreated dishes. Neurospheres were then cultured in CDM with FGF-8 (#423-F8, R&D Systems) for 7 days. On day 12, the medium was supplemented with Purmorphamine (#540220-5MG, EMD Millipore) and FGF-8 for midbrain precursor patterning. From day 19, hMOs were embedded in Matrigel, with the addition of BDNF (#450-02, PeproTech) and GDNF (#450-10, PeproTech) until their four-month maturation.

### Nicotinamide Riboside Treatment

The hMOs were subjected to daily treatment with a 1 mM concentration of NR for a duration of two months. The NR was provided by Evandro Fei Fang from the University of Oslo.

### hMO Staining Protocol

hMO cells were spread onto cover slides and allowed to air dry at ambient temperature. Post-drying, they were fixed with 4% PFA (#28908, Thermo Scientific). This was followed by two rounds of PBS rinsing, after which they were submerged in a 20% sucrose-PBS mix and kept overnight at 4°C. The next day, cells were blocked for 2 hours at room temperature, then incubated overnight at 4°C with primary antibodies, anti-SOX2 (Mouse, #ab171380, Abcam), anti-PAX6 (Rabbit, #ab5790, Abcam), anti-MAP2 (Chicken, #ab5392, Abcam). After a comprehensive 3-hour PBS wash, cells were exposed overnight to secondary antibodies in a chilled, moisture-retentive, light-shielded environment. Coverslips were then mounted using Fluor mount-G with DAPI (#0100-20, SouthenBiotech).

### Imaging & Analysis Techniques

Images were obtained through a Leica TCS SP8 confocal laser scanning apparatus, with the assistance of Leica LAS X software. Fiji (Fiji Is Just ImageJ) served as our primary tool for image adjustments and evaluations. For quantitative analysis, we randomly picked 6-10 sectors. Fluorescence intensity measurements required us to convert individual channel images to 8-bit format using Image J. To standardize thresholding across images, we adhered to default settings. The chosen algorithm and parameters were also set to default. Fluorescence intensity was gauged via the mean gray value (Mean), with the equation: Mean = IntDen/area. All data interpretations and illustrations were facilitated by GraphPad Prism 8.0.2 (GraphPad Software, Inc).

### Single-cell RNA Sequencing (scRNA-seq) and Data Analysis

#### Organoid Dissociation and Single Cell Isolation

To collect the organoids, they were removed from the culture medium and washed with 1x PBS (Invitrogen, #10010-23). The organoids were then finely cut into pieces of 1-2 mm using ophthalmic scissors. Next, the organoid pieces were digested in 2 ml of CellLive™ Tissue Dissociation Solution (#1190062, Singleron Biotechnologies,) at 37°C for 15 minutes in a 15-ml centrifuge tube (#62.5544.003, Sarstedt,), with continuous agitation on a thermal shaker. The degree of dissociation was periodically checked under a light microscope. Following digestion, the suspension was filtered through a 40-µm sterile strainer (Greiner#542040). The cells were then centrifuged at 350 x g for 5 minutes at 4°C, and the resulting cell pellets were resuspended in 1 ml of PBS. To assess cell viability and count, the cells were stained with a 0.4% w/v solution of Trypan Blue (#15250-061, Gibco). The cell number and viability were determined using a hemocytometer under a light microscope.

#### Data Preprocessing Steps

Fastq data preprocessing was achieved using CeleScope® (v.1.14.1; www.github.com/singleron-RD/CeleScope; Singleron Biotechnologies GmbH), keeping default settings. After discarding low-quality reads, we utilized STAR (https://github.com/alexdobin/STAR) to map sequences to the human GRCh38 reference. Gene annotation was based on Ensembl 92, and reads were assigned to genes with feature count (https://subread.sourceforge.net/). Cell identification leveraged a negative bimodal distribution, resulting in a count matrix detailing the UMI per gene in every cell. For further analysis, we employed the scanpy package (29) in Python and Seurat library [2] in R. QC metrics were extracted from the gene count matrix, eliminating non-viable cells and potential doublets, as well as discarding cellular debris.

#### Clustering & Cell Annotation

Integration of data was handled by Harmany (30) within the scanpy workflow. Clustering analysis was executed on a combined dataset from four samples (control 1, control 2, disease, treated) via Scanpy. A neighborhood graph was formulated, followed by unsupervised cell clustering using the Leiden algorithm. Based on known markers sourced from various studies, clusters were annotated. dopamine (DA)-1/2 was annotated since it expressed a well-known DA neuronal marker NR4A2 as well as DCX and SYT1 among others (31, 32). Using cell typists (33) and human brain models (34), ventral midbrain neurons (VMN) were identified. Fig. 1E tabulates gene expressions relevant to cell cluster annotations. Control samples were formulated by merging Control 1 (Detroit 551) and Control 2 (CCD-1079Sk), retaining original cell annotations.

**Figure 1.**
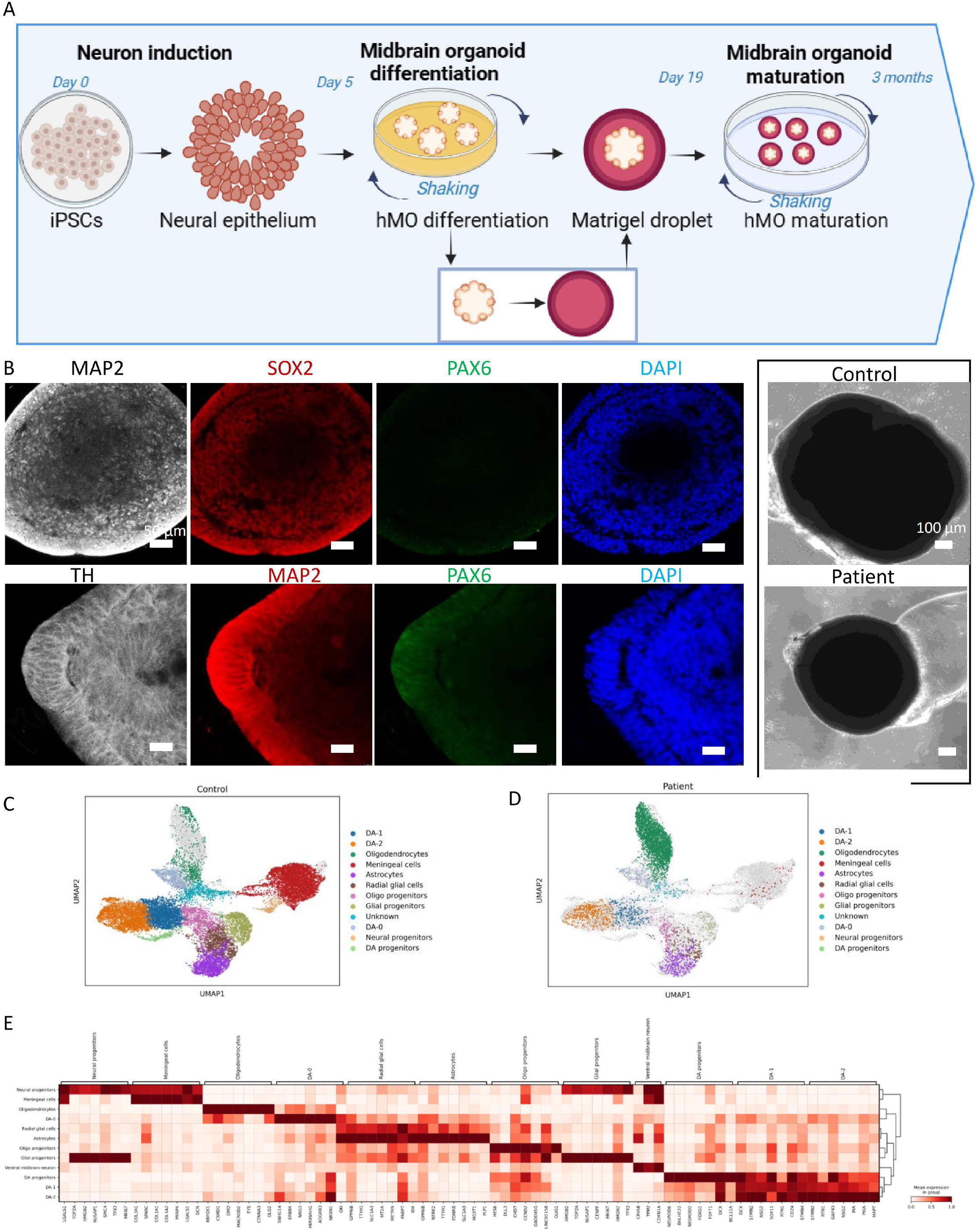
Development of hMOs from iPSCs. A: Diagrammatic illustration of the differentiation process from iPSCs to hMOs. Illustration created using BioRender.com. B: Fluorescence microscopy images showcasing control and POLG iPSC-derived hMOs at 3 months, marked with antibodies against TH, SOX2, PAX6, and MAP2. Cell nuclei are counterstained with DAPI (blue). Accompanying phase-contrast images depict hMOs cultivated from control and POLG patient iPSCs. The scale bar represents 50 µm. C-D: Uniform Manifold Approximation and Projection (UMAP) for cell annotation in hMOs derived from control (C) and POLG patients (D). E: Differential expression profiles across various cell clusters identified within hMOs.

#### Differential Expression and Pathway Analysis

We conducted a differential expression analysis utilizing *<u>the scanpy.tl.rank_genes_groups</u>* function, focusing on the distinguished cell clusters and various cell types. We computed both the proportions of cells based on samples and the absolute counts for individual clusters within subgroups. For custom groupings of clusters, we employed the Wilcoxon rank sum test and applied the Benjamini-Hochberg method for p-value correction during the differential expression analysis.

#### Enrichment Analysis

We conducted enrichment analyses for Gene Ontology (GO) and the Kyoto Encyclopedia of Genes and Genomes (KEGG) by closely examining genes labeled as upregulated and downregulated. For these analyses, we utilized an online tool found at https://www.bioinformatics.com.cn, as previously referenced (35). This in-depth analysis helps pinpoint and comprehend notable gene and pathway enrichments, providing greater insight into their significance and interplay within the biological framework.

## Results

### Generating 3D POLG-hMOs

Using dual-SMAD inhibition combined with FGF-8b and the Sonic hedgehog (SHH) agonist PM, (28) (Figure 1A). To guide neuroectoderm differentiation towards the floor plate, iPSCs were first differentiated to neuroepithelium (neural rosettes), using dual SMAD inhibition. Next, these were transitioned to suspension culture, via dissociation to produce 3D neurospheres, which were maintained on an orbital shaker. The neurospheres were then caudalized for a week with FGF-8b, followed by ventralization with FGF-8b and PM for an additional week. Next, the midbrain organoids were encapsulated in Matrigel droplets and matured using BDNF and GDNF. We performed the differentiation in both human iPSC lines harbouring a POLG mutation and 2 control iPSCs, Detroit 551 and CCD-1079Sk lines. The derived hMOs were maintained in suspension on an orbital shaker, for over 2 months for subsequent evaluation. By 3 months, it was observed that POLG hMOs were smaller compared to the control (Fig. 1B).

SOX2 is a conserved transcription factor in vertebrates, involved in central nervous system development (36). In our midbrain organoids, SOX2 displayed positive staining, especially in younger organoids. Concurrently, the mature neuronal marker microtubule-associated protein 2 (MAP2) also demonstrated positive staining, hinting at neuronal formation (Fig. 1B). In 60-day-old midbrain samples, we observed both SOX2 and MAP2 expression by immunostaining, which was predominantly on the neuroepithelium’s apical surface (Fig. 1B). Notably, the hMOs lacked PAX6 expression, which is a favourable sign of midbrain specification, PAX6 is essential for forebrain development (37).

These data suggest that by utilizing our protocol, we successfully generated 3D POLG-hMOs from iPSCs, resulting in hMOs that, after three months. Interestingly we observed a smaller diameter in POLG-derived hMOs as compared to controls. Notably, the hMOs displayed positive staining for the transcription factor SOX2 and the neuronal marker MAP2, while the absence of Pax6 expression suggested that the hMOs had a midbrain identity.

### Alterations in Single-Cell Transcriptomic Profiles in POLG Patient-Derived hMOs

To provide deeper insight into the transcriptomic changes occurring at the single-cell level, we conducted single-cell RNA sequencing (scRNA-seq) analysis. Our analysis revealed that both control and POLG patient-derived organoids encompassed distinct cell populations, including DA0 cells, DA1 cells, DA2 cells, oligodendrocytes, radial glial cells, oligodendrocyte progenitors, glial progenitors, ventral midbrain neurons, astrocytes, neural progenitors, DA progenitors, and meningeal cells (Fig. 1C and D), identified based on their gene expression patterns (Fig. 1E).

In our prior research, we observed a significant reduction in DA neurons (21). Delving deeper, we categorized these DA neurons into three specific sub-populations, using their distinct gene expression profiles, as described earlier (38). We labelled these as intermediate cells in the dopaminergic lineage DA0, along with two subtypes of dopaminergic neurons, DA1 and DA2. In POLG patient-derived hMOs, we observed a significant reduction in DA1 neurons (4.88%), DA2 neurons (10.36%), and ventral midbrain neurons (VMN, 1.97%) (Fig. 2B), as compared to control, where DA1 (13.83%), DA2 (16.40%), and VMN (4.00%) (Fig. 2A) were slightly higher. However, there was a marginal decrease in DA0 neurons (3.24%) and DA progenitors (0.02%) in POLG hMOs (Fig. 2B) compared to control (Fig. 2A), where DA0 (3.97%) and DA progenitors (1.47%) exhibited slightly higher levels.

**Figure 2.**
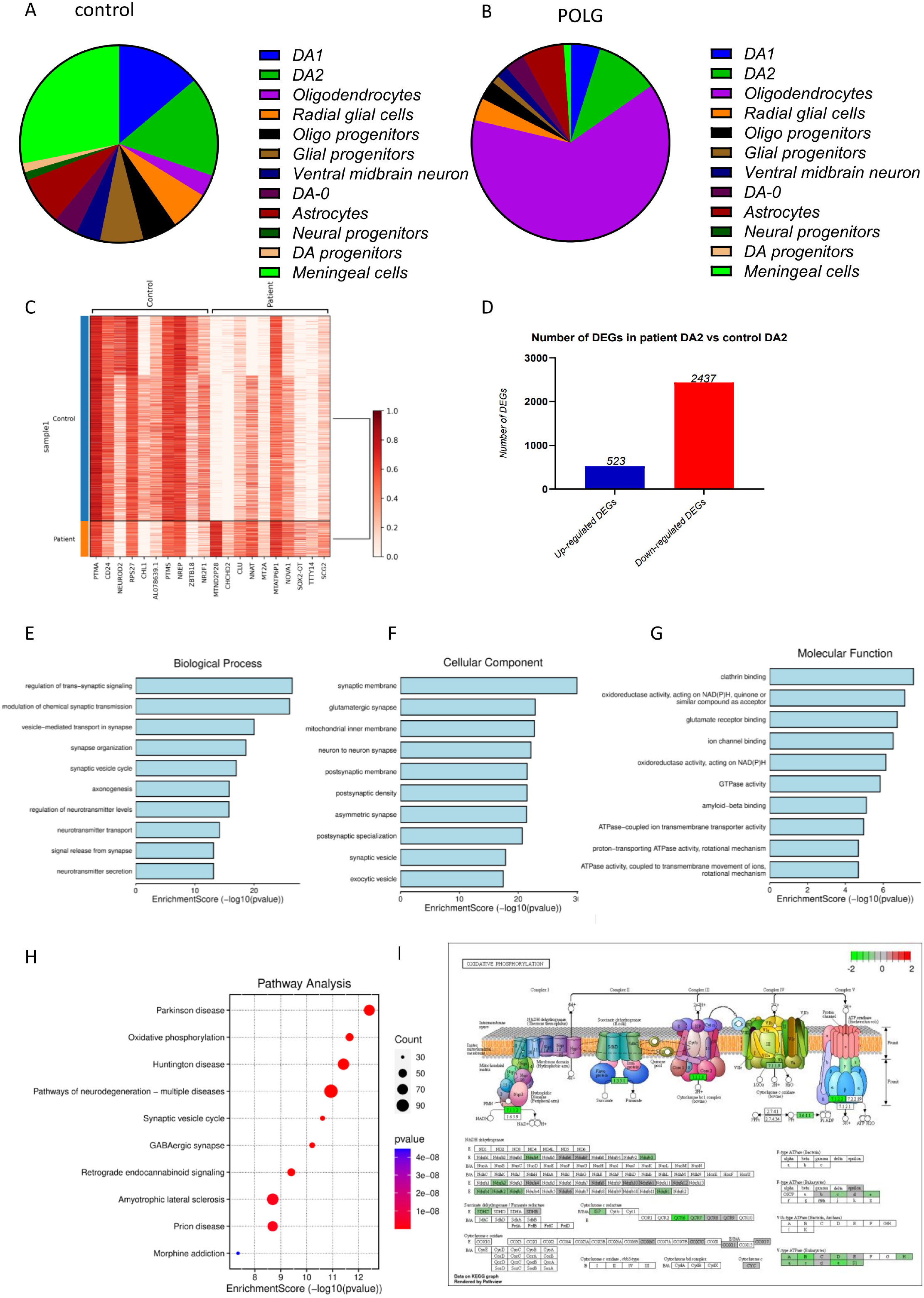
scRNA-seq Analysis of hMOs Derived from Control and POLG Patients. A-B: Pie charts depicting the distribution of predominant cell clusters, identified by specific marker expression, in hMOs from controls (A) and POLG patients (B). C: Heatmap illustrating the differential expression of genes in DA2 cluster cells from POLG patient-derived hMOs compared to those from controls, with marker genes along the x-axis and sample identifiers on the y-axis. D: Histogram detailing the counts of DEGs, both upregulated and downregulated, in DA2 cluster cells from control and POLG patient-derived hMOs. E: Bar chart of the top ten GO BP terms associated with downregulated DEGs in the hMOs of POLG patients relative to those from controls. F: Bar chart showing the top ten GP CC terms linked to downregulated DEGs in the hMOs of POLG patients compared to control-derived hMOs. G: Bar chart presenting the top ten GO MF terms enriched in downregulated DEGs in the hMOs of POLG patients versus those from controls. H: Scatter plot demonstrating the KEGG pathway enrichment for downregulated DEGs in DA2 cluster cells of hMOs from POLG patients, with the enrichment score calculated as −log10 (P-value) and counts indicating the number of DEGs associated with each Pathway ID. I: Schematic and comprehensive analysis of the downregulated DEGs in the oxidative phosphorylation pathway in the hMOs of POLG patients versus those from controls.

In the context of gene expression analysis, the patient-derived hMOs’ DA2 cells showed a significant downregulation as compared to controls (Fig. 2C). Specifically, we identified 523 differentially expressed genes (DEGs) that were upregulated and 2437 that were downregulated (Fig. 2D).

This data identified diverse cell populations in both control and POLG patient-derived organoids. Notably, POLG hMOs exhibited a significant reduction in specific DA neuron subtypes, including DA1 and DA2 cells, as well as VMN. Gene expression analysis within the DA2 cell population revealed extensive downregulation of genes in POLG hMOs, providing insights into the molecular changes associated with neurological pathology.

### Functional Insights from Gene Ontology (GO) and KEGG Enrichment Analysis in POLG hMOs DA2 Neurons

Exploration of GO enrichment for the downregulated DEGs within DA2 cells revealed several potentially dysregulated biological processes (BP). These included the regulation of trans-synaptic signalling, modulation of chemical synaptic transmission, vesicle-mediated transport in synapses, synapse organization, synaptic vesicle cycle, axonogenesis, regulation of neurotransmitter levels, neurotransmitter transport, signal release from synapses, and neurotransmitter secretion (Fig. 2E).

Regarding cellular components (CC), the top ten GO terms encompassed synaptic membrane, glutamatergic synapse, mitochondrial inner membrane, neuron-to-neuron synapse, postsynaptic membrane, postsynaptic density, asymmetric synapse, postsynaptic specialization, synaptic vesicle, and exocytic vesicle (Fig. 2F).

Furthermore, molecular function (MF) enrichment analysis identified the top ten terms related to clathrin binding, oxidoreductase activity (acting on NAD(P)H, quinone, or similar compounds as acceptors), glutamate receptor binding, ion channel binding, oxidoreductase activity (acting on NAD(P)H), GTPase activity, amyloid-beta binding, ATPase-coupled ion transmembrane transporter activity, proton-transporting ATPase activity (rotational mechanism), and ATPase regulator activity (Fig. 2G).

KEGG pathway analysis illuminated a propensity towards neurodegenerative pathways in patient-derived DA2 neurons. Examination of the downregulated DEGs’ KEGG pathways highlighted Parkinson’s, Huntington’s, and ALS, with notable associations with oxidative phosphorylation, retrograde endocannabinoid signalling, GABAergic synapse, and the synaptic vesicle cycle (Fig. 2H). Notably, the “Oxidative Phosphorylation” pathway emerged as a central hub (Fig. 2I), with downregulated DEGs predominantly clustered within mitochondrial respiratory chain complexes (Fig. 2J). Specifically, genes from complexes I and IV (*NDUFS3, NDUFS2, NDUFS7, NDUFS6, NDUFA8, NDUFB6, NDUFS4, NDUFB3, NDUFA5, NDUFA2, NDUFA4, NDUFB2, NDUFV3, NDUFC1, NDUFB1, ATP6V1E1, ATP6V1C1, ATP6V0D1, ATP6V0A1, ATP6V1H, ATP6V0B, ATP6V1D, ATP6V0E2, ATP6V1A, ATP5F1D, ATP5MD, ATP5F1E, ATP5PD, ATP5PB, ATP5MPL, ATP5MC1, ATP5MC3, ATP5ME, ATP5IF1, CYC1*) and a single gene from complex II (*SDHB*), along with multiple genes from complexes III (*UQCRH, UQCRHL, UQCR10, UQCRFS1, UQCRQ) and IV (COX11, COX7A2L, COX6C*), underscored profound mitochondrial disturbances in patient-derived DA2 neurons compared to controls (Fig. 2I).

These findings provide comprehensive insights into the transcriptional alterations in POLG patient-derived hMOs and shed light on critical pathways and processes implicated in neurodegenerative disorders.

### Mitochondrial and Neural-Related Transcriptomic Alterations in POLG hMOs DA2 Neurons

The pathway analysis of downregulated DEGs within POLG hMOs’ DA2 neurons uncovered substantial perturbations, particularly in mitochondrial and neural-related processes. Within the mitochondrial domain, an array of critical functions and processes were downregulated in POLG hMOs DA2 neurons. These included mitochondrial translational elongation, mitochondrial translational termination, mitochondrial respiratory chain complex assembly, oxidative phosphorylation, mitochondrial translation, ATP metabolic processes, mitochondrial transport, mitochondrial gene expression, respiratory electron transport chain, macro-autophagy, and NADH dehydrogenase complex assembly (Fig. 3A).

**Figure 3.**
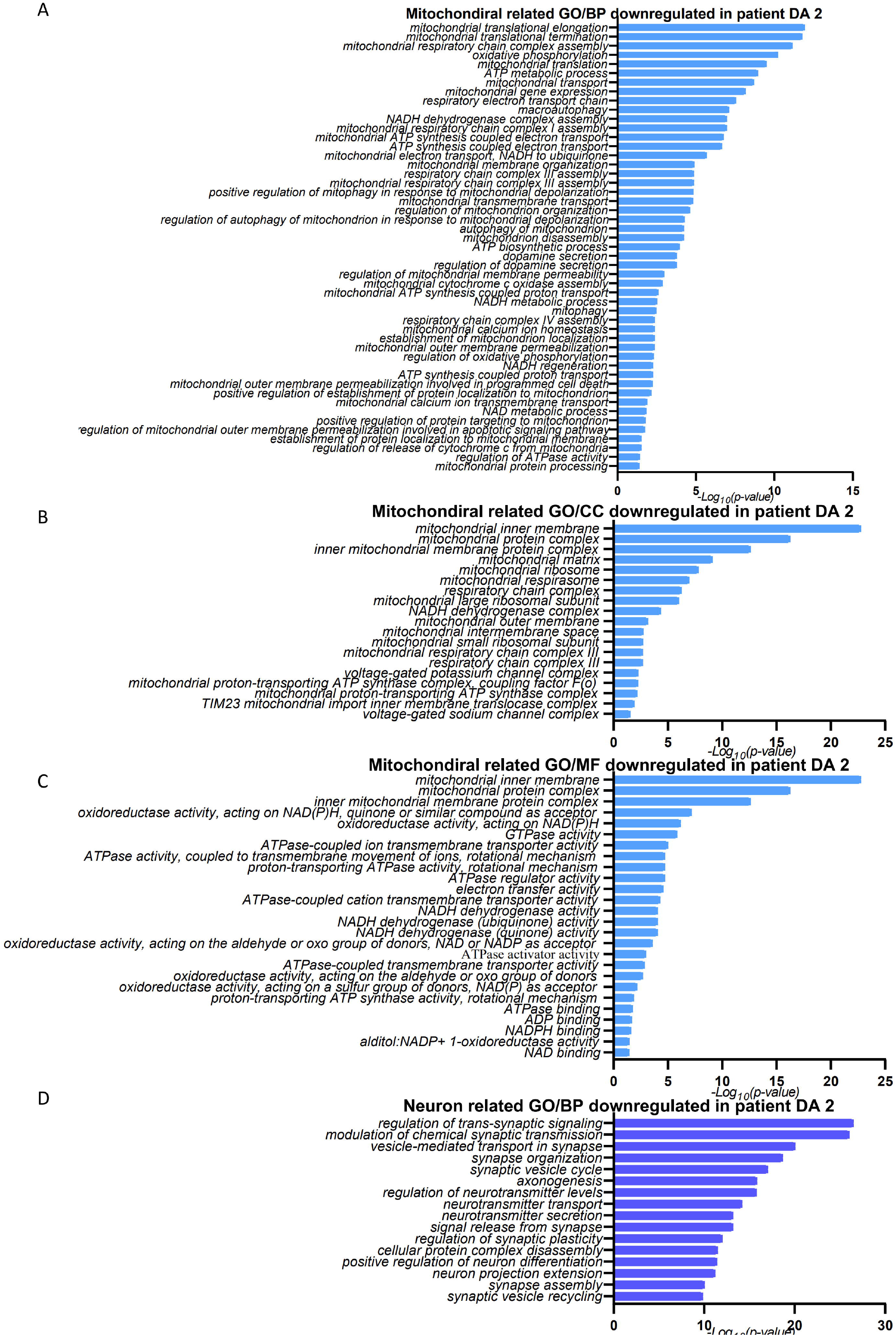
Downregulation of Mitochondrial and Neuronal GO Terms and KEGG Pathways in DA2 Neurons of POLG Patient-Derived hMOs. A: Downregulated GO BP terms related to mitochondria in DA2 neurons from POLG patient-derived hMOs. B: Downregulated GO CC terms associated with mitochondrial functions in DA2 neurons of hMOs from POLG patients. C: Downregulated GO MF terms pertaining to mitochondrial activities in DA2 neurons of POLG patient hMOs. D: Principal neural-related GO BP terms that are downregulated in DA2 neurons of hMOs from POLG patients.

Furthermore, we observed notable decreases in MFs associated with oxidoreductase activities, including those acting on NAD(P)H and quinone compounds. Additionally, there was downregulation in ATPase activities coupled to ion transmembrane movement, electron transfer, and proton transmembrane transport. The electron transfer activities reflect compromised mitochondrial electron transport processes (Fig. 3B and C). These findings underscore the impact of *POLG* mutations on mitochondrial function.

In the context of neural-related processes, our analysis revealed downregulation in critical functions such as the regulation of trans-synaptic signalling, modulation of chemical synaptic transmission, vesicle-mediated transport in synapses, synapse organization, synaptic vesicle cycle, axonogenesis, regulation of neurotransmitter levels, neurotransmitter transport, neurotransmitter secretion, signal release from synapses, and regulation of synaptic plasticity (Fig. 3D). These changes indicate compromised neuronal communication and synaptic function within POLG hMOs DA2 neurons.

This analysis underscores the interconnection between mitochondrial and neural-related alterations in POLG hMOs DA2 neurons, shedding light on the molecular underpinnings of POLG-related neurodegenerative pathology.

### Insights into Gene Expression Alterations in POLG hMOs VMNs

The gene expression profiles of POLG hMOs’ ventral midbrain neurons (VMNs) revealed a substantial downregulation compared to control cells (Fig. 4A). KEGG pathway analysis unveiled a notable inclination towards pathways associated with neurodegenerative processes in patient-derived VMNs. The downregulated DEGs were linked to pathways encompassing ribosome, coronavirus disease (COVID-19), glycolysis/gluconeogenesis, biosynthesis of amino acids, and carbon metabolism (Fig. 4B).

**Figure 4.**
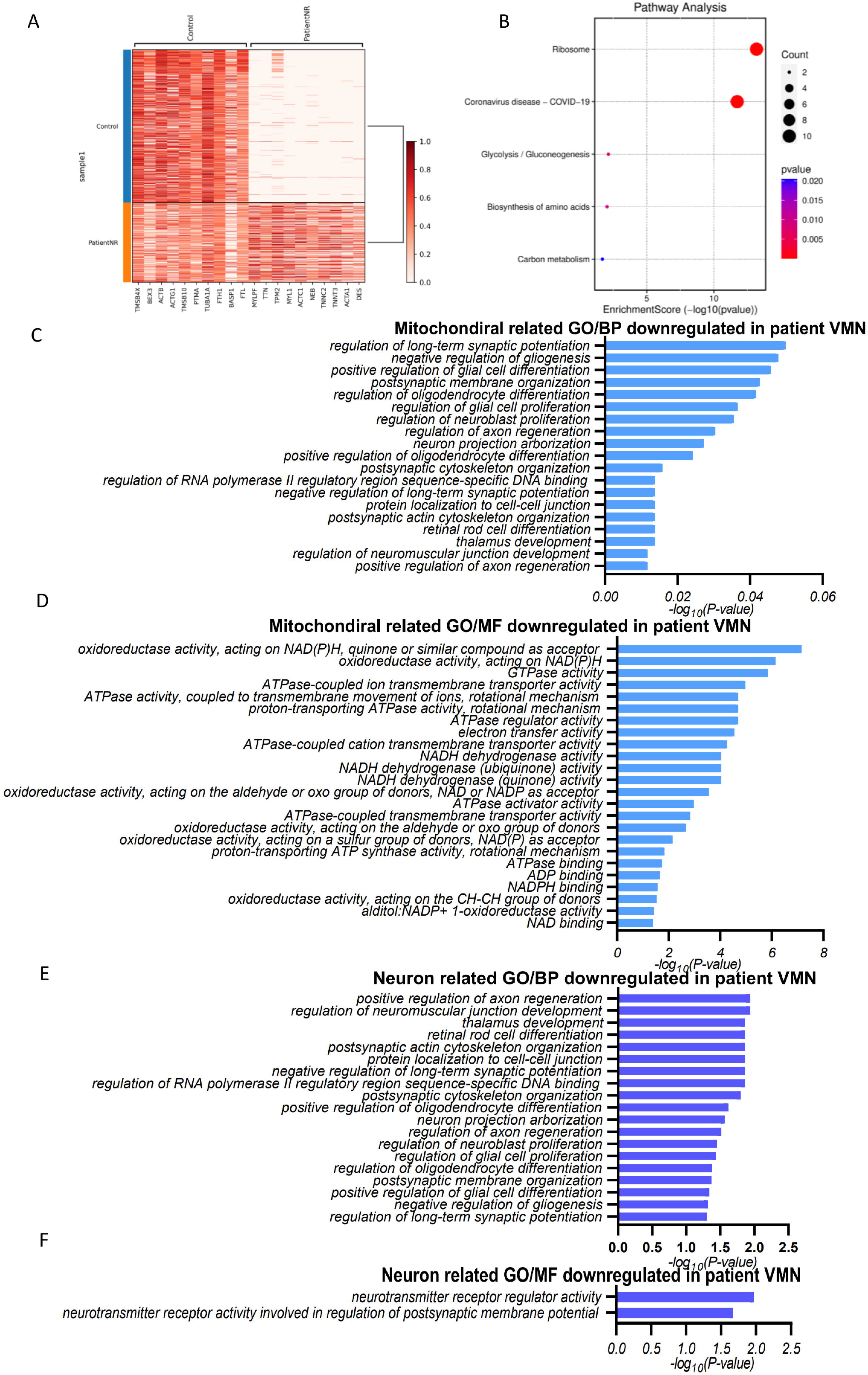
Downregulation of Mitochondrial and Neuronal GO Terms and KEGG Pathways in VMN of hMOs Derived from POLG Patients. A: Heatmap illustrating the differential expression of genes in VMN cluster cells from POLG patient-derived hMOs compared to those from controls, with marker genes along the x-axis and sample identifiers on the y-axis. B: A scatter plot reveals the KEGG pathway enrichment within the VMN, where the enrichment score is calculated as the negative logarithm (base 10) of the P-value, and the counts correspond to the number of DEGs associated with each pathway. C: Mitochondrial-associated BP terms from the GO database, which are found to be downregulated in the VMNs of hMOs from POLG patients, are listed. D: Mitochondrial-associated MF terms from the GO database, showing downregulation in the VMNs of POLG patient-derived hMOs, are detailed. E: The most significantly downregulated neural-associated BP terms from the GO database in the VMNs of hMOs from POLG patients are highlighted. F: The top-downregulated neural-associated MF terms from the GO database in the VMNs of POLG patient hMOs are presented.

Within the mitochondrial-related BP highlighted by GO enrichment for downregulated DEGs, we observed processes related to positive regulation of axon regeneration, regulation of neuromuscular junction development, thalamus development, retinal rod cell differentiation, postsynaptic actin cytoskeleton organization, protein localization to cell-cell junctions, negative regulation of long-term synaptic potentiation, regulation of RNA polymerase II regulatory region sequence-specific DNA binding, postsynaptic cytoskeleton organization, positive regulation of oligodendrocyte differentiation, neuron projection arborization, and more (Fig. 4C).

Regarding mitochondrial-related CC, the top GO terms included oxidoreductase activity (acting on NAD(P)H, quinone, or similar compounds as acceptors), GTPase activity, ATPase-coupled ion transmembrane transporter activity, proton-transporting ATPase activity (rotational mechanism), ATPase regulator activity, electron transfer activity, ATPase-coupled cation transmembrane transporter activity, NADH dehydrogenase activity, NADH dehydrogenase (ubiquinone) activity, NADH dehydrogenase (quinone) activity, and many others (Fig. 4D).

Similarly, GO enrichment for downregulated DEGs within VMNs unveiled neural-related BP. These encompassed functions like positive regulation of axon regeneration, regulation of neuromuscular junction development, thalamus development, retinal rod cell differentiation, postsynaptic actin cytoskeleton organization, protein localization to cell-cell junctions, negative regulation of long-term synaptic potentiation, regulation of RNA polymerase II regulatory region sequence-specific DNA binding, postsynaptic cytoskeleton organization, positive regulation of oligodendrocyte differentiation, neuron projection arborization, regulation of neuroblast proliferation, and more (Fig. 4E). Additionally, GO enrichment for downregulated DEGs within VMNs also revealed neural-related MF. These included functions such as neurotransmitter receptor regulator activity and neurotransmitter receptor activity involved in regulating postsynaptic membrane potential (Fig. 4F).

Overall, this analysis sheds light on the gene expression alterations observed in POLG hMOs VMNs, as well as the molecular changes associated with these neurodegenerative conditions.

### Nicotinamide Riboside Treatment Enhanced Transcriptomic Changes in NR-Treated POLG hMOs DA Neurons

To gain a further understanding of the transcriptomic alterations at the single-cell level, we subjected POLG patient-derived hMOs to NR treatment, combined with scRNA-seq analysis on DA (DA2) and VMNs. This revealed that NR-treated hMOs maintained the same cell populations as untreated POLG organoids but displayed increased numbers of DA neurons and VMNs (Fig. 5A). Interestingly, VMNs exhibited a significant increase (from 1.97% to 5.59%) in NR-treated patient-derived hMOs, however, we did not observe a comparable rise in DA2 neurons (10.37%) (Fig. 5B).

**Figure 5.**
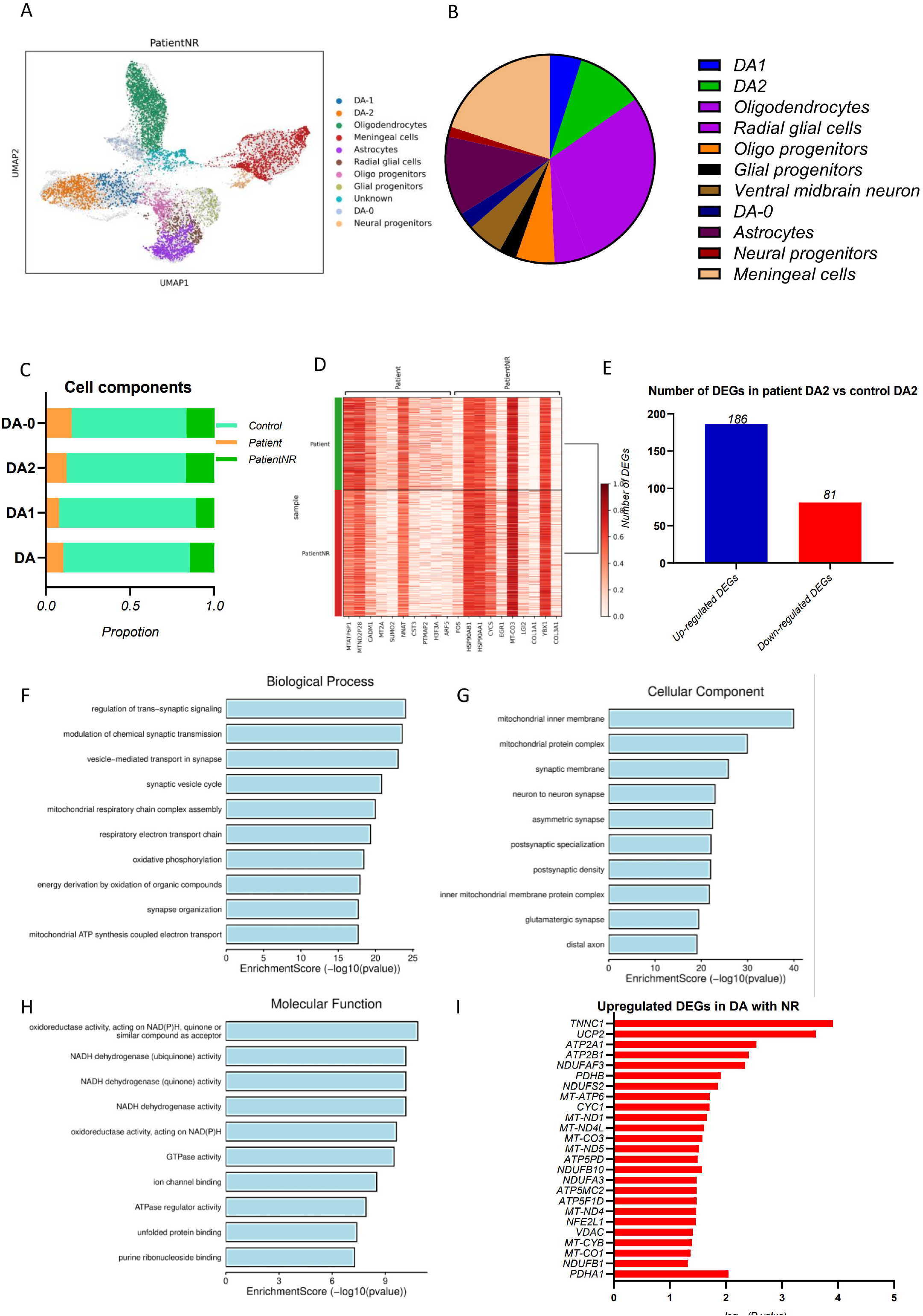
Sc RNA-seq analysis in control and POLG patient hMO treated with NR. A: UMAP analysis for cell annotation in control and POLG patient-derived hMOs with nr treatment. B: Pie charts comparing major cell cluster proportions in POLG patient-derived hMOs with and without NR treatment, based on specific marker expression. C: Comparison of different DA cell cluster proportions in control hMOs, POLG patient-derived hMOs and POLG patient-derived hMOs treated with NR. D: Heatmap of gene expression in DA neurons from POLG patient-derived hMOs with NR treatment versus untreated, with markers along the x-axis and sample identifiers on the y-axis. E: Histogram of upregulated and downregulated DEGs in DA neurons from POLG patient-derived hMOs with NR treatment versus untreated samples. F: Top ten downregulated GO BP terms in DA neurons of hMOs. G: Top ten downregulated GO CC terms in DA neurons from POLG patient-derived hMOs with NR treatment compared to untreated samples. H: Top ten downregulated GO MF terms in DA neurons from POLG patient-derived hMOs with nr treatment compared to untreated samples. I: Top ten upregulated mitochondrial-related genes in DA neurons from POLG patient-derived hMOs with NR treatment versus untreated samples.

Next, we performed gene expression analysis, the NR-treated patient hMOs’ DA cells, and a significant upregulation was observed compared to untreated samples (Fig. 5E). Specifically, 186 DEGs were upregulated and 81 DEGs downregulated (Fig. 5E). GO enrichment for the upregulated DEGs within DA cells revealed prominent BP. These included the regulation of trans-synaptic signalling, modulation of chemical synaptic transmission, vesicle-mediated transport in synapses, synaptic vesicle cycle, mitochondrial respiratory chain complex assembly, respiratory electron transport chain, oxidative phosphorylation, energy derivation by oxidation of organic compounds, synapse organization, and mitochondrial ATP synthesis coupled electron transport (Fig. 5F).

Examining the CC highlighted the top ten GO terms, which included mitochondrial inner membrane, mitochondrial protein complex, synaptic membrane, neuron-to-neuron synapse, asymmetric synapse, postsynaptic specialization, postsynaptic density, inner mitochondrial membrane protein complex, glutamatergic synapse, and distal axon (Fig. 5G).

MF enrichment analysis identified the top ten terms, which included oxidoreductase activity acting on NAD(P)H, quinone, or similar compounds as acceptors, NADH dehydrogenase activity, NADH dehydrogenase (ubiquinone) activity, NADH dehydrogenase (quinone) activity, oxidoreductase activity acting on NAD(P)H, GTPase activity, ion channel binding, ATPase regulator activity, unfolded protein binding, and purine ribonucleoside binding (Fig. 5H).

Lastly, NR-treated patient-derived DA neurons showed the downregulated DEGs identified specific genes, including *NDUFB1, MT-CO1, MT-CYB, VDAC, NFE2L1, MT-ND4, ATP5F1D, ATP5MC2, NDUFA3, NDUFB10, ATP5PD, MT-ND5, MT-CO3, MT-ND4L, MT-ND1, CYC1, MT-ATP6, NDUFS2, PDHB, PDHA1, NDUFAF3, ATP2B1, ATP2A1, UCP2,* and *TNNC1,* which were downregulated post-NR treatment (Fig. 5I).

This data suggests that NR treatment of POLG patient-derived hMOs led to increased neurons, with upregulation in gene expression in both related to synaptic function and mitochondrial activity in DA cells, indicating potential therapeutic benefits for POLG-related neurodegeneration.

### Nicotinamide Riboside Treatment Ameliorates the Mitochondrial and Neural-Related Transcriptomic Alterations observed in POLG hMOs-derived DA Neurons

The transcriptomic responses observed in POLG hMOs’ DA neurons post-NR treatment identified perturbations, especially in mitochondrial and neural-related processes. When we examined the effects of NR treatment on mitochondrial BP in POLG hMOs DA neurons, downregulation was evident. Specifically, the highlighted processes included mitochondrial respiratory chain complex assembly, respiratory electron transport chain, oxidative phosphorylation, mitochondrial ATP synthesis coupled electron transport, cellular respiration, mitochondrial translational elongation, and more (Fig. 6A). Our CC analysis for mitochondrial processes revealed notable increases in structures such as mitochondrial inner membrane, mitochondrial protein complex, inner mitochondrial membrane protein complex, mitochondrial respirasome, and others (Fig. 6B). Further the MF for mitochondrial processes, upregulation in activities like oxidoreductase activity, NADH dehydrogenase activity, proton transmembrane transporter activity, and electron transfer activity among others was observed (Fig. 6C).

**Figure 6:**
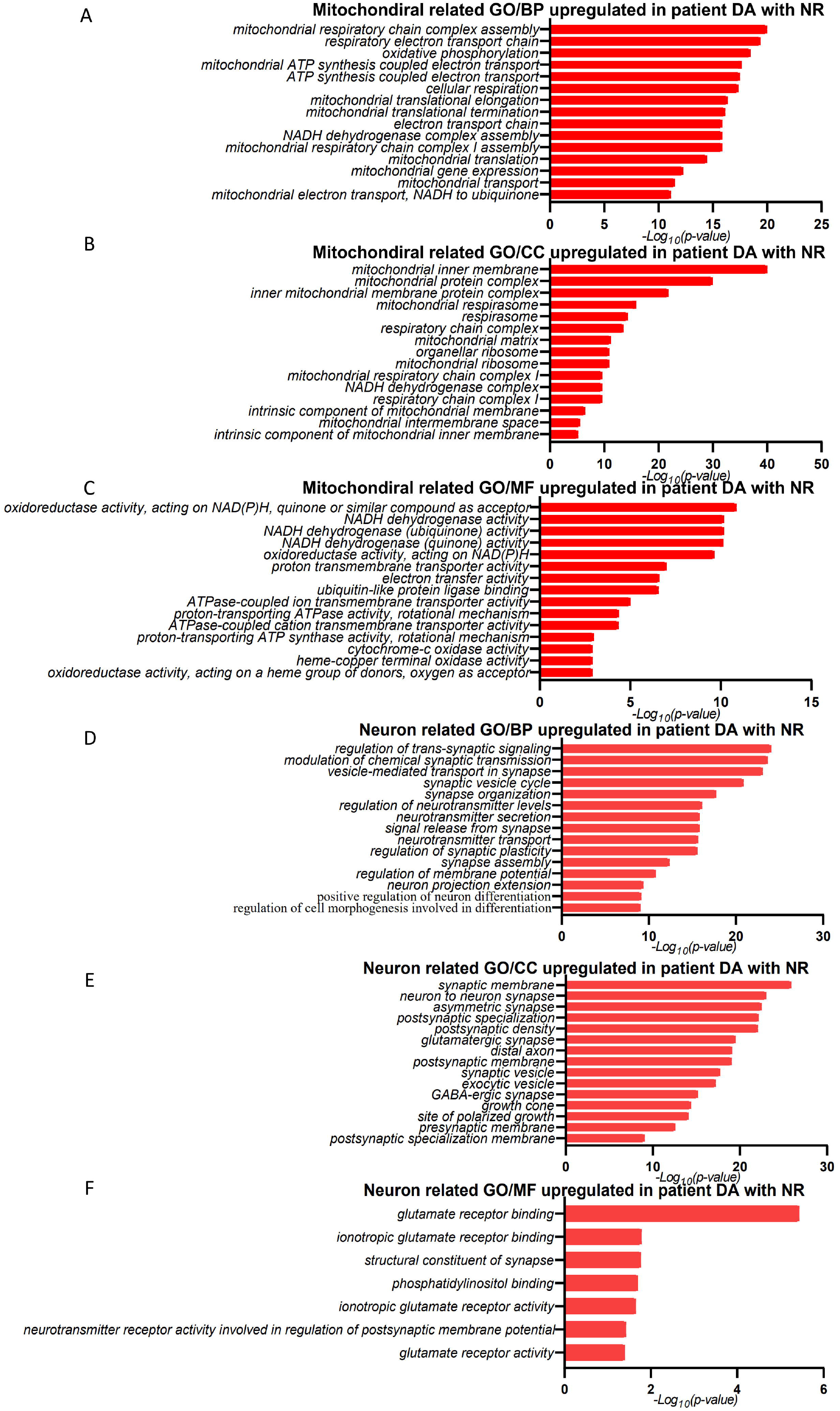
Upregulation of Mitochondrial and Neuronal Pathways in DA from hMOs of POLG Patients Following NR Treatment. A-C: Indications of upregulated mitochondrial-related GO terms in the BP, CC, and MF categories in DA in POLG patient-derived hMOs post-NR administration. D-F: Display of enhanced neuronal-related GO terms in the BP, CC, and MF categories in DA in POLG patient-derived hMOs post-NR administration.

Turning our attention to neural-related processes, we found upregulation in BP including regulation of trans-synaptic signalling, modulation of chemical synaptic transmission, vesicle-mediated transport in the synapse, and more (Fig. 6D). Analyzing the CC associated with neural processes, there were increases in elements such as synaptic membrane, neuron to neuron synapse, asymmetric synapse, and postsynaptic density to name a few (Fig. 6E). Lastly, our investigation into the MF pertaining to neural processes revealed upregulated activities, highlighting roles such as synaptic membrane activity, neuron to neuron synapse function, and asymmetric synapse activity among others (Fig. 6F).

In this analysis of the transcriptomic changes in POLG hMOs’ DA neurons following NR treatment, we detected changes. Mitochondrial processes were upregulated, with key functions like oxidative phosphorylation and cellular respiration affected, mitochondrial structures increased, and a rise in specific mitochondrial functions. Concurrently, neural-related processes showed an upsurge, particularly in synaptic activities and components.

### Nicotinamide Riboside Treatment Improves Transcriptomic Alterations in POLG hMOs DA2 Neurons

In the context of gene expression analysis within the NR-treated POLG hMOs’ DA2 cells, upregulation was observed compared to controls (Fig. 7A). Specifically, we identified 147 DEGs that were upregulated and 79 DEGs that were downregulated (Fig. 7B).

**Figure 7:**
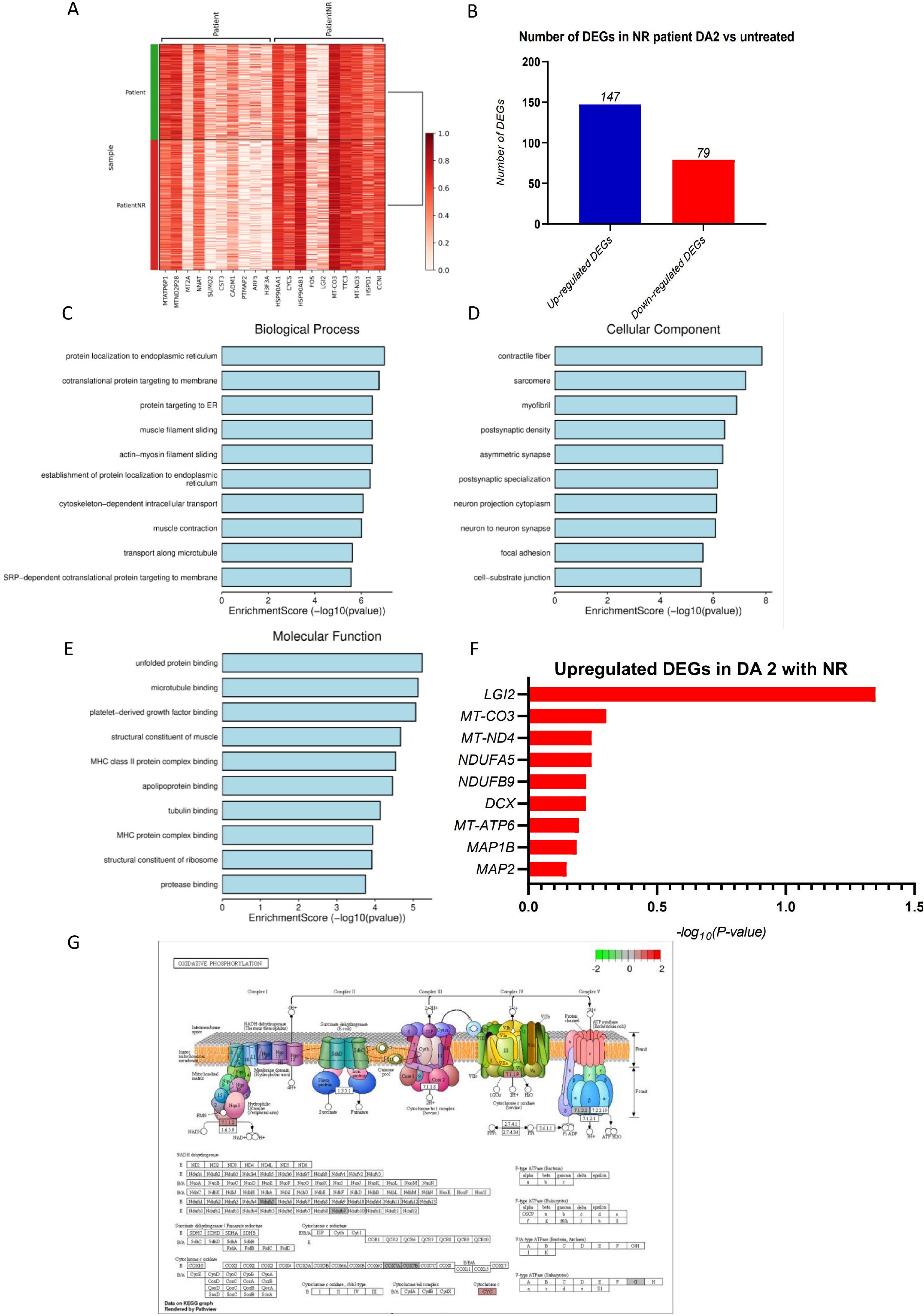
Single-cell RNA-Seq Analysis Showcasing Mitochondrial Dysfunction in POLG patient-derived hMOs with NR Treatment. A: Heatmap showing the upregulated and downregulated DEGs in DA2 from POLG-derived hMOs following NR administration compared to untreated POLG-derived hMOs, with gene markers along the x-axis and individual sample identifiers along the y-axis. B: Bar graph showing number fo the upregulated and downregulated DEGs s when comparing NR-treated DA2 patient samples to untreated POLG-derived hMOs. C: Top ten upregulated GO BP terms in DA2 from POLG-derived hMOs following NR administration. D: Top ten upregulated GO CC terms in DA2 from POLG-derived hMOs following NR administration. E: Top ten upregulated related GO MF terms showing downregulation DA2 from POLG-derived hMOs following NR administration. F: Upregulated mitochondrial related genes in DA2 from POLG-derived hMOs following NR administration. G: Upregulated genes linked to oxidative phosphorylation that are influenced by NR treatment in DA2 from POLG-derived hMOs following NR administration.

GO enrichment for the upregulated DEGs within DA2 cells, highlighted processes which included protein localization to the endoplasmic reticulum, cotranslational protein targeting to the membrane, protein targeting to ER, muscle filament sliding, actin-myosin filament sliding, establishment of protein localization to endoplasmic reticulum, cytoskeleton-dependent intracellular transport, and muscle contraction, with further notable processes like transport along microtubule and SRP-dependent cotranslational protein targeting to the membrane (Fig. 7C).

In terms of CC, the top GO terms encapsulated contractile fibre, sarcomere, myofibril, postsynaptic density, asymmetric synapse, postsynaptic specialization, neuron projection cytoplasm, and neuron-to-neuron synapse. Additionally, notable structures were focal adhesion and cell-substrate junction (Fig. 7D).

Moving onto MF enrichment, the leading terms observed related to unfolded protein binding, microtubule binding, platelet-derived growth factor binding, structural constituent of muscle, MHC class II protein complex binding, apolipoprotein binding, tubulin binding, and MHC protein complex binding. We also observed structural constituents of ribosome and protease binding as significant functions (Fig. 7E). We found that NR-treated patient-derived DA neurons showed the upregulated DEGs identified specific genes, including mitochondrial related genes (*LGI2, MT-CO3, MT-ND4, NDUFA5, NDUFB9, MT-ATP6*), and neural related genes (*DCX, MAP1B* and *MAP2*), where were mitochondrial and neural related genes (Fig. 7F). We detected the upregulated DEGs predominantly clustered within mitochondrial respiratory chain complexes after NR treatment, including the genes from complexes I and IV (*NDUFA5, NDUFB9, COX7A* and *COX7B*), underscored the improvement of the mitochondrial disturbances in NR-treated patient-derived DA2 neurons compared to untreated samples (Fig. 7G).

In summary, in NR-treated POLG hMOs’ DA2 cells, 147 genes were upregulated and 79 were downregulated compared to controls. The GO enrichment analysis highlighted key processes in protein localization, synapse structures, and protein binding functions.

### Nicotinamide Riboside Treatment Improves the Transcriptomic Alterations in POLG hMOs DA2 Neurons

Our analysis of GO enrichment for the upregulated DEGs within NR-treated POLG hMOs’ DA2 neurons, we identified key BP related to mitochondria. These processes encompassed the establishment of protein localization to the endoplasmic reticulum, establishment of mitochondrion localization, microtubule-mediated mitochondrion transport, oxidative phosphorylation, ATP metabolic process, and electron transport chain. Further significant processes included mitochondrial electron transport, NADH dehydrogenase complex assembly, and cellular respiration, among others (Fig. 8A).

**Figure 8:**
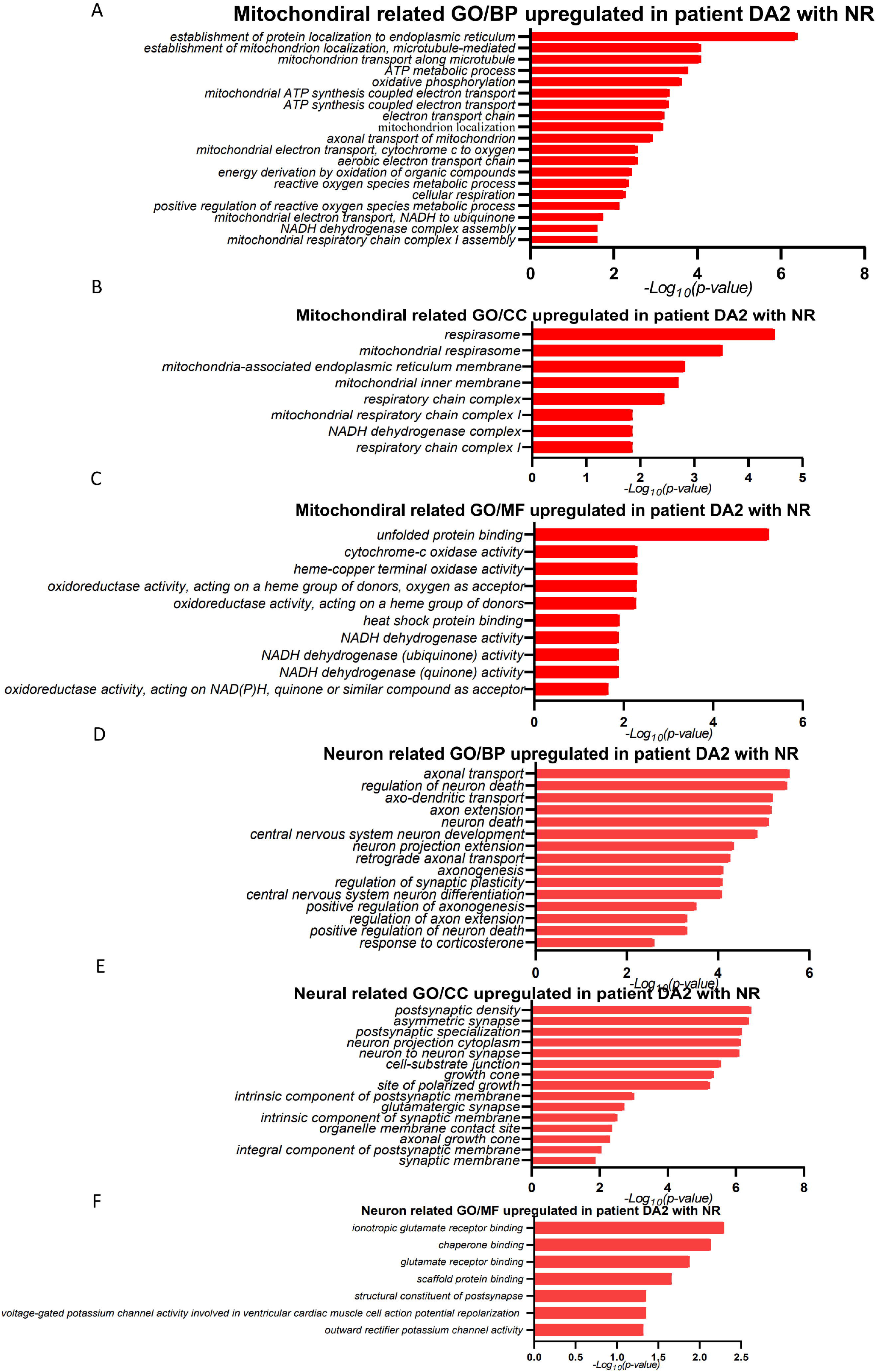
Enhancement of Mitochondrial and Neuronal Functions in POLG-patient DA2 neurons in POLG-derived hMOs with NR Administration. A-c: Representations of upregulated mitochondrial related GO terms across BP, CC, and MF DA2 from POLG-derived hMOs following NR administration. D-F: Representations of upregulated neuronal related GO terms across BP, CC, and DA2 from POLG-derived hMOs following NR administration.

Regarding the CC associated with mitochondria, our findings highlighted terms such as respirasome, mitochondrial inner membrane, respiratory chain complex I, NADH dehydrogenase complex, and mitochondria-associated endoplasmic reticulum membrane. Other notable components identified include the mitochondrial respiratory chain complex and respiratory chain complex I (Fig. 8B).

For the MF enrichment, the top terms related to mitochondrial function included unfolded protein binding, cytochrome-c oxidase activity, oxidoreductase activity, heat shock protein binding, and NADH dehydrogenase activity. Additionally, we identified other functions like oxidoreductase activity, acting on NAD(P)H, quinone or similar compounds as acceptors (Fig. 8C).

In a neural context, the dominant BP identified included axonal transport, regulation of neuron death, axon extension, neuron death, and central nervous system neuron development. Additional processes of interest were neuron projection extension, axonogenesis, and regulation of synaptic plasticity (Fig. 8D). While neural-related CCs showed postsynaptic density, asymmetric synapse, postsynaptic specialization, neuron projection cytoplasm, and neuron-to-neuron synapse. We also observed other structures such as the growth cone, synaptic membrane, and glutamatergic synapse (Fig. 8E).

Neural-related MF enrichment, our top findings included ionotropic glutamate receptor binding, chaperone binding, glutamate receptor binding, and scaffold protein binding. Other significant functions were related to the structural constituent of post-synapse and voltage-gated potassium channel activity (Fig. 8F).

Overall, the analysis of upregulated DEGs in NR-treated POLG hMOs’ DA2 neurons uncovered key mitochondrial and neural processes, components, and functions. Including oxidative phosphorylation and electron transport in a mitochondrial context, while neural findings included axonal transport, neuron development, and postsynaptic structures and activities.

### Nicotinamide Riboside Treatment Improves the Transcriptomic Alterations observed in POLG hMOs VMNs

Gene expression analysis of NR-treated POLG hMOs’ derived VMNs, showed significant upregulation as compared to untreated (Fig. 9A). Specifically, 310 DEGs that were upregulated while only 2 DEGs that were downregulated (Fig. 9B). GO enrichment showed that the upregulated DEGs in NR treated VMNs, the top BPs included cotranslational protein targeting to membrane, SRP-dependent cotranslational protein targeting to membrane, nonsense-mediated decay of nuclear-transcribed mRNA, protein targeting to ER, and establishment of protein localization to endoplasmic reticulum. Additionally, translational initiation, protein localization to endoplasmic reticulum, viral transcription, and viral gene expression, among others (Fig. 9C).

**Figure 9:**
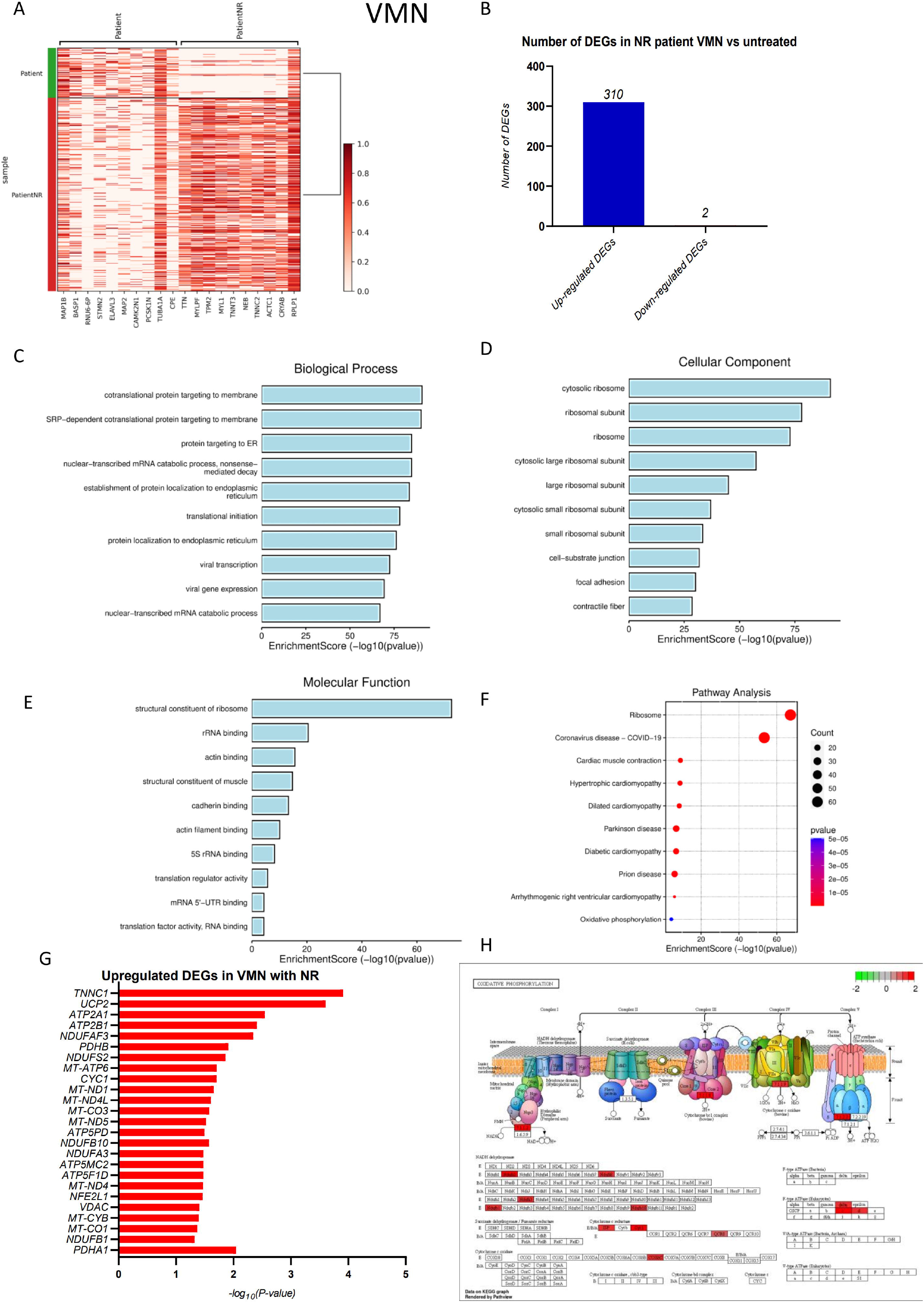
scRNA-Seq Reveals Mitochondrial and Pathway Alterations in VMN in POLG-derived hMOs with NR Treatment. A: Gene expression profiles in VMN in POLG-derived hMOs post-NR, plotted with gene markers versus sample labels. B: Histogram illustrating the count of DEGs in NR-treated VMN in POLG-derived hMOs post-NR versus untreated counterparts. C-F: Downregulated mitochondrial-related GO terms and KEGG pathway enrichment analysis in VMN from POLG-derived hMOs, with a detailed look at the DEGs and their associated pathways post-NR treatment. G: Upregulated mitochondrial KEGG pathway terms related to DEGs in VMN from POLG-derived hMOs post-NR treatment. H: Graphical analysis detailing the upregulated genes in oxidative phosphorylation pathways in VMN from POLG-derived hMOs post-NR treatment.

For CCs, top GO terms included cytosolic ribosome, ribosomal subunits, ribosome, and large and small ribosomal subunits. As well as cell-substrate junction, focal adhesion, and contractile fiber (Fig. 9D).

MF enrichment highlighted prominent terms such as structural constituent of ribosome, rRNA binding, actin binding, structural constituent of muscle, and cadherin binding. Further, actin filament binding, 5S rRNA binding, translation regulator activity, mRNA 5’-UTR binding, and translation factor activity relating to RNA binding as significant functions (Fig. 9E).

KEGG pathway analysis indicated a trend towards neurodegenerative pathways in NR-treated patient-derived VMNs. Our review of the upregulated DEGs’ KEGG pathways pinpointed areas such as Ribosome, Coronavirus disease - COVID-19, Cardiac muscle contraction, and Hypertrophic cardiomyopathy, among others, also Parkinson’s disease, prion disease, and Oxidative Phosphorylation (Fig. 9F). The “Oxidative Phosphorylation” pathway stood out (Fig. 2I), with upregulated DEGs predominantly clustered within mitochondrial respiratory chain complexes (Figure 9G). Specifically, genes from various complexes, including those from *NDUFB1, MT-CO1, MT-CYB, VDAC, NFE2L1, MT-ND4, ATP5F1D, ATP5MC2, NDUFA3, NDUFB10, ATP5PD, MT-ND5, MT-CO3, MT-ND4L, MT-ND1, CYC1, MT-ATP6, NDUFS2, PDHB, PDHA1, NDUFAF3, ATP2B1, ATP2A1, UCP2,* and *TNNC1*, highlighted significant mitochondrial anomalies in NR treated patient-derived VMN neurons when juxtaposed with untreated samples (Fig. 9H).

We next looked at Further GO enrichment for the upregulated DEGs within NR-treated POLG hMOs’ VMNs, we identified key BP related to mitochondria. These processes encompassed cellular response to oxygen levels, mitochondrial ATP synthesis coupled electron transport, ATP synthesis coupled proton transport, and cellular response to hypoxia. Further significant processes involved in mitochondrial function included regulation of transcription in response to hypoxia, NADH dehydrogenase complex assembly, aerobic respiration, and mitochondrial transport, among others (Fig. 10A).

**Figure 10:**
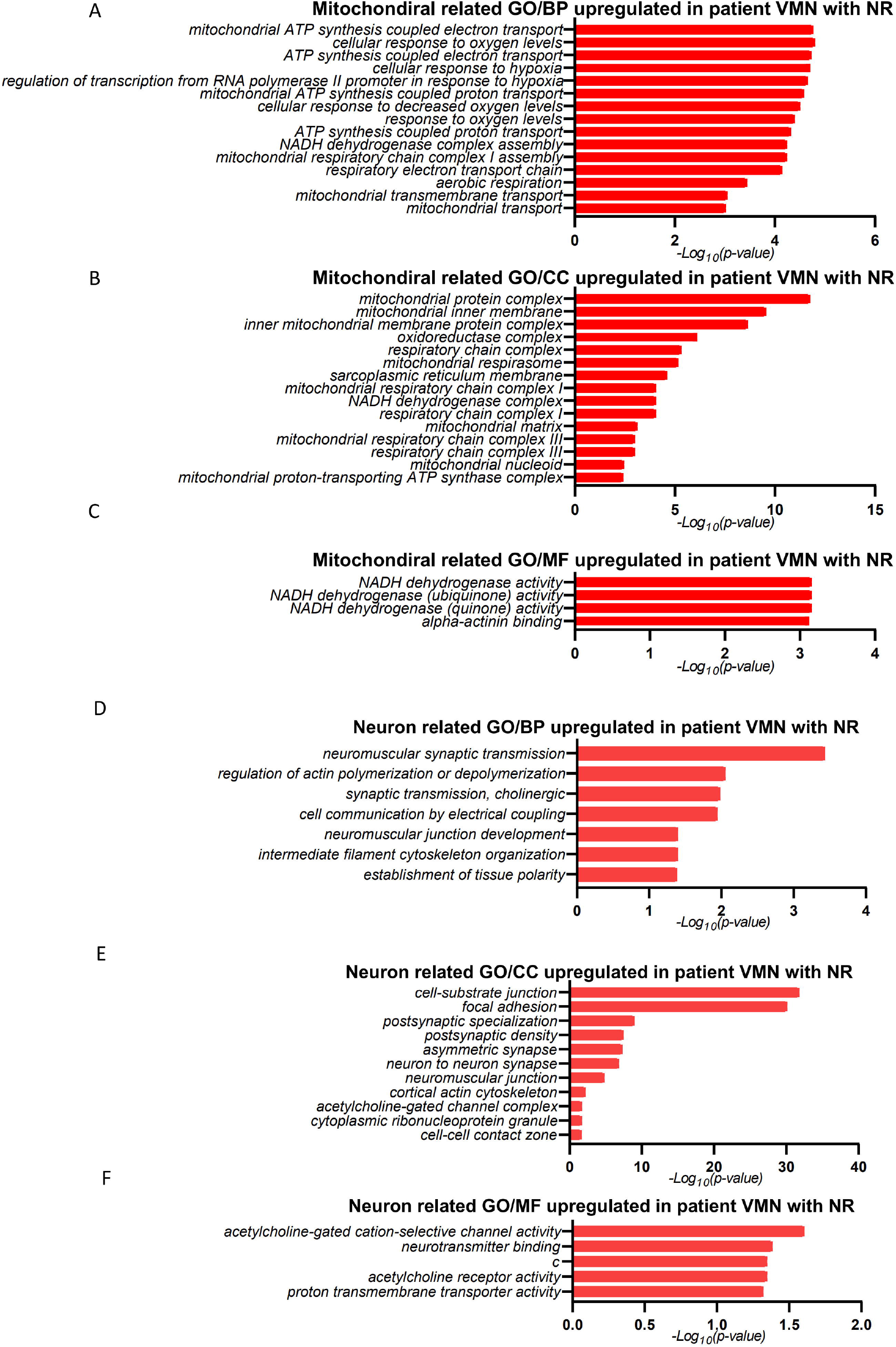
Upregulation of Mitochondrial and Neuronal Pathways in VMN HMOs of POLG Patients Following NR Treatment. A-C: Indications of upregulated mitochondrial-related GO terms in the BP, CC, and MF categories in VMN in hMOs post-NR administration. D-F: Display of enhanced neuronal-related GO terms in the BP, CC, and MF categories in VMN in hMOs post-NR administration.

Regarding the CC associated with mitochondria, our findings emphasized terms such as mitochondrial protein complex, mitochondrial inner membrane, oxidoreductase complex, and mitochondrial respirasome. Other notable components spotlighted were the sarcoplasmic reticulum membrane, mitochondrial matrix, and mitochondrial proton-transporting ATP synthase complex (Fig. 10B).

For the MF enrichment, the top terms related to mitochondrial function included NADH dehydrogenase activities, with various substrates, and alpha-actinin binding (Fig. 10C).

In the neural context, the dominant BP identified included neuromuscular synaptic transmission, synaptic transmission in cholinergic systems, neuromuscular junction development, and establishment of tissue polarity. Additional neural processes spotlighted were the regulation of actin polymerization or depolymerization and intermediate filament cytoskeleton organization (Fig. 10D).

In terms of neural-related CC, our highlighted terms were cell-substrate junction, focal adhesion, postsynaptic specialization, and neuron-to-neuron synapse. We also observed other structures such as neuromuscular junction, cortical actin cytoskeleton, and acetylcholine-gated channel complex (Fig. 10E).

For neural-related MF enrichment, our top findings showcased terms like acetylcholine-gated cation-selective channel activity, neurotransmitter binding, and acetylcholine receptor activity (Fig. 10F).

Overall, this analysis of upregulated DEGs in NR-treated POLG hMOs’ VMN neurons illuminated central mitochondrial and neural processes, components, and functions. Mitochondrial highlights spanned areas of ATP synthesis and response to oxygen levels, while neural discoveries concentrated on synaptic transmission, neuromuscular development, and acetylcholine activities.

## Discussion

Our previous study (39), describes the generation of midbrain-specific organoids from both embryonic stem cells and iPSCs. This methodology produced organoids with midbrain characteristics via expression of midbrain markers, while largely excluding forebrain and hindbrain indicators. Of critical importance was the observation that these organoids exhibited the presence of dopaminergic neurons characteristic of the midbrain, a significant implication for studying disorders like PD and their association with mitochondrial dysfunction. Building on these findings, our current study harnesses this organoid model to further explore the impact of *POLG* mutations. This research underscores how these mutations may impinge on transcriptomic changes in hMOs. Moreover, this study reveals the therapeutic potential of NR, an NAD^+^ precursor, in counteracting the adverse effects of *POLG* mutations.

The expression of SOX2, a transcription factor paramount for CNS development, along with MAP2, suggests that the generated hMOs had the capacity for neuronal differentiation. This was further corroborated by the absence of PAX6, which emphasizes the fidelity of our differentiation protocol in creating midbrain-specific neurons. The dual expression of SOX2 and MAP2 on the apical surface of the neuroepithelium provides additional affirmation of the neuronal differentiation trajectory.

scRNA-seq investigations unveiled the distinct cellular repertoire in both control and POLG-derived hMOs. The reduced DA neuron subpopulations in POLG hMOs echoes the findings of (40, 41), providing a molecular basis for the noted vulnerability of these neurons in POLG-associated disorders. Furthermore, the pronounced downregulation of mitochondrial and neural-related pathways within the DA2 cell population broadens our molecular understanding, illuminating the intricate regulatory changes stemming from *POLG* mutations.

Intriguingly, the functional insights derived from the GO and KEGG enrichment analyses suggest a significant impact on pathways and processes that are pivotal for neuronal function and health. The perturbations observed in pathways like oxidative phosphorylation and others associated with neurodegeneration point to mitochondrial disturbances. The interconnectedness of mitochondrial and neural-related transcriptional alterations underscores the potential of mitochondrial health in neuronal homeostasis (42, 43).

Our exploration of VMNs showed downregulation in POLG hMOs, revealing potential underpinnings of the neurodegenerative phenotype in POLG-related disorders. The inclination towards neurodegenerative processes and pathways could provide a basis for the phenotypic manifestations observed clinically in these disorders. Our data indicates that disruptions in mitochondrial function might be the epicenter of the symptoms characterizing POLG mutations.

One of this study’s prime findings is the substantial and multifaceted impact of *POLG* mutations on mitochondrial function. Mitochondria, often regarded as cellular powerhouses, are indispensable for neuronal function given the high energy demands of neurons. Therefore, any disturbance in mitochondrial function could very well be the linchpin in POLG-related neurodegeneration. Moreover, the neural-related transcriptional changes we observed, especially those associated with synapses and neurotransmitter systems, could explain the neurological deficits commonly associated with POLG disorders.

To decipher the underlying transcriptomic shifts, we treated POLG patient-derived hMOs with NR and analyzed the neuronal subtypes DA (DA2) and VMNs. Interestingly, NR treatment, akin to the observations by Parker et al. (2019), appeared to have restorative effects on the cellular populations, with a noticeable surge in VMNs. The transcriptomic responses post-NR treatment, particularly within the DA2 neurons and VMNs, further emphasize the modulatory potential of mitochondrial and neural processes in response to this treatment.

Upon closer inspection of the gene expression data from NR-treated patient hMOs’ DA cells, significant upregulation in gene expression was observed compared to untreated samples. To that end GO enrichment analysis revealed upregulation in biological processes related to trans-synaptic signaling, synaptic vesicle cycles, and mitochondrial respiratory chain complex assembly, among others. Additionally, the cellular components and molecular functions were prominently related to mitochondrial inner membranes and oxidoreductase activity. Further examination of the transcriptomic responses post-NR treatment highlighted significant changes in mitochondrial and neural-related processes. Mitochondrial biological processes showed a noticeable downregulation, while the cellular components for these processes, like the mitochondrial inner membrane, showed notable increases. Also, neural-related processes demonstrated an upsurge, particularly in synaptic activities and components post-NR treatment. The DA2 cells of the NR-treated POLG hMOs, showed a significant upregulation in gene expression, with GO enrichment for upregulated DEGs revealing prominent biological processes related to protein localization to the endoplasmic reticulum, muscle contraction, and microtubule-based transport. On the cellular and molecular front, we observed structures related to contractile fibres and functions like unfolded protein binding.

When considering the VMNs of NR-treated POLG hMOs, a substantial upregulation in gene expression was observed. The primary biological processes covered protein targeting the ER and viral gene expression. The cellular components were mostly centred on ribosomal structures, while molecular functions primarily involved structural constituents of ribosomes and actin binding. Additionally, KEGG pathway analysis indicated potential links to neurodegenerative pathways in the NR-treated VMNs. Notably, the “Oxidative Phosphorylation” pathway was prominent, with upregulated DEGs clustered within mitochondrial respiratory chain complexes. On deeper GO enrichment analysis for the upregulated DEGs in NR-treated VMNs showed key mitochondrial-related biological processes, including cellular responses to oxygen levels and ATP synthesis. The cellular components and molecular functions were predominantly related to mitochondrial structures and NADH dehydrogenase activities. On the neural front, processes like neuron death and axonogenesis were dominant.

In summary, this study, supplemented by insights from contemporary literature, provides insights into *POLG* mutations and their ramifications on neuronal health. It underscores the promise of therapeutic avenues that emphasize mitochondrial restoration, offering hope for interventions in POLG-related disorders.

## Conclusion

This comprehensive study of 3D iPSC-derived POLG hMOs provides insight into their differentiation and the molecular underpinning of POLG-related neurodegenerative disorders. The described hMO protocol yields midbrain-centric organoids produces the desired markers and lacks unnecessary ones. Of note, was the diminished growth of POLG hMOs potentially pointing to challenges arising from *POLG* mutations.

The expressions of SOX2 and MAP2 in the organoids confirmed their ability to differentiate into neurons. scRNA-seq. evaluation also highlights the vulnerability of DA neuron subgroups in POLG-related ailments and highlights considerable gene expression shifts in the DA2 cell cluster. The findings from GO and KEGG enrichment analyses further emphasized the potential role of mitochondrial health and its interplay with neural functions, reinforcing the notion that mitochondrial disruption is central to POLG-associated neurodegeneration. Moreover, our observations underscored the significance of mitochondrial activities for neurons, which have substantial energy requirements. The noted alterations in neural transcriptions could potentially be the root cause of the frequently observed neurological issues in POLG disorders.

Furthermore, NR treatment of POLG patient-derived hMOs leads to significant transcriptional changes with respect to both DA neurons and VMNs. NR-treated organoids retained most cell populations, there was a marked increment in VMNs, more so in DA2. The upregulated gene expression post-NR treatment, especially in the mitochondrial and neural domains, could have therapeutic implications for POLG-related disorders. Notably, the shifts in processes associated with the endoplasmic reticulum and ribosomal structures also warrant attention for potential therapeutic avenues.

In essence, our study robustly delineates the multifaceted effects of *POLG* mutations on both mitochondrial and neuronal functions. These findings could pave the way for deeper insights into POLG-related neurodegenerative disorders and offer avenues for therapeutic interventions.

## Data Availability

The RNA sequencing analysis read count data can be accessed in the NCBI Gene Expression Omnibus (GEO) data deposit system with an accession number GSE241743. All other data are available from the corresponding author upon request.

## Conflict of Interest

All authors declare that the research was conducted in the absence of any commercial or financial relationships that could be construed as a potential conflict of interest.

## Funding

This work was supported by the following funding: K.L was supported by the University of Bergen Meltzers Høyskolefonds (#103517133) and Gerda Meyer Nyquist Legat (#103816102).

## Author’s contributions

K.L contribute to the conceptualization; A.C and T. Y contribute to the methodology and the investigation; K.L, G.J. S and A.C contribute to the writing original draft; all authors contribute to the writing review and editing; K.L contribute to the funding acquisition and to the resources; K.L contributes to the supervision. All authors agree to the authorships.

## Acknowledgements

Our gratitude goes to Evandro Fei Fang of the University of Oslo for generously supplying the NR. We also extend our appreciation to the teams at the Molecular Imaging Centre Core Facility in the University of Bergen for their invaluable expertise and support in confocal imaging recording.

## Notes

### Competing Interest Statement

The authors have declared no competing interest.

## Reference

1. Sato T, Imaizumi K, Watanabe H, Ishikawa M, Okano H. Generation of region-specific and high-purity neurons from human feeder-free iPSCs. Neuroscience Letters. 2021;746:135676.

2. Corti S, Faravelli I, Cardano M, Conti L. Human pluripotent stem cells as tools for neurodegenerative and neurodevelopmental disease modeling and drug discovery. Expert Opin Drug Discov. 2015;10(6):615–29.

3. Nguyen KV, Østergaard E, Ravn SH, Balslev T, Danielsen ER, Vardag A, et al. POLG mutations in Alpers syndrome. Neurology. 2005;65(9):1493–5.

4. Bhattacharya A, Choi WWY, Muffat J, Li Y. Modeling Developmental Brain Diseases Using Human Pluripotent Stem Cells-Derived Brain Organoids - Progress and Perspective. J Mol Biol. 2022;434(3):167386.

5. Qian X, Song H, Ming GL. Brain organoids: advances, applications and challenges. Development. 2019;146(8).

6. Sun AX, Ng HH, Tan EK. Translational potential of human brain organoids. Ann Clin Transl Neurol. 2018;5(2):226–35.

7. Kofman S, Mohan N, Sun X, Ibric L, Piermarini E, Qiang L. Human mini brains and spinal cords in a dish: Modeling strategies, current challenges, and prospective advances. J Tissue Eng. 2022;13:20417314221113391.

8. Shariati L, Esmaeili Y, Haghjooy Javanmard S, Bidram E, Amini A. Organoid technology: Current standing and future perspectives. Stem Cells. 2021;39(12):1625–49.

9. Choi J, Kim S, Jung J, Lim Y, Kang K, Park S, et al. Wnt5a-mediating neurogenesis of human adipose tissue-derived stem cells in a 3D microfluidic cell culture system. Biomaterials. 2011;32(29):7013–22.

10. Sunildutt N, Parihar P, Chethikkattuveli Salih AR, Lee SH, Choi KH. Revolutionizing drug development: harnessing the potential of organ-on-chip technology for disease modeling and drug discovery. Front Pharmacol. 2023;14:1139229.

11. Svrakic DM, Zorumski CF. Neuroscience of Object Relations in Health and Disorder: A Proposal for an Integrative Model. Front Psychol. 2021;12:583743.

12. Lancaster MA, Renner M, Martin CA, Wenzel D, Bicknell LS, Hurles ME, et al. Cerebral organoids model human brain development and microcephaly. Nature. 2013;501(7467):373–9.

13. Paşca AM, Sloan SA, Clarke LE, Tian Y, Makinson CD, Huber N, et al. Functional cortical neurons and astrocytes from human pluripotent stem cells in 3D culture. Nat Methods. 2015;12(7):671–8.

14. Haycock JW. 3D cell culture: a review of current approaches and techniques. Methods Mol Biol. 2011;695:1–15.

15. D’Avanzo F, Rigon L, Zanetti A, Tomanin R. Mucopolysaccharidosis Type II: One Hundred Years of Research, Diagnosis, and Treatment. Int J Mol Sci. 2020;21(4).

16. Seidel K, Siswanto S, Brunt ER, den Dunnen W, Korf HW, Rüb U. Brain pathology of spinocerebellar ataxias. Acta Neuropathol. 2012;124(1):1–21.

17. Gomez-Giro G, Arias-Fuenzalida J, Jarazo J, Zeuschner D, Ali M, Possemis N, et al. Synapse alterations precede neuronal damage and storage pathology in a human cerebral organoid model of CLN3-juvenile neuronal ceroid lipofuscinosis. Acta Neuropathologica Communications. 2019;7(1):222.

18. Smits LM, Reinhardt L, Reinhardt P, Glatza M, Monzel AS, Stanslowsky N, et al. Modeling Parkinson’s disease in midbrain-like organoids. NPJ Parkinsons Dis. 2019;5:5.

19. Choi SH, Kim YH, Hebisch M, Sliwinski C, Lee S, D’Avanzo C, et al. A three-dimensional human neural cell culture model of Alzheimer’s disease. Nature. 2014;515(7526):274–8.

20. Liang KX, Kristiansen CK, Mostafavi S, Vatne GH, Zantingh GA, Kianian A, et al. Disease-specific phenotypes in iPSC-derived neural stem cells with POLG mutations. EMBO Mol Med. 2020;12(10):e12146.

21. Liang KX, Vatne GH, Kristiansen CK, Ievglevskyi O, Kondratskaya E, Glover JC, et al. N-acetylcysteine amide ameliorates mitochondrial dysfunction and reduces oxidative stress in hiPSC-derived dopaminergic neurons with POLG mutation. Experimental Neurology. 2021;337:113536.

22. Chen A, Kristiansen CK, Hong Y, Kianian A, Fang EF, Sullivan GJ, et al. Nicotinamide Riboside and Metformin Ameliorate Mitophagy Defect in Induced Pluripotent Stem Cell-Derived Astrocytes With POLG Mutations. Front Cell Dev Biol. 2021;9:737304.

23. Kristiansen CK, Chen A, Høyland LE, Ziegler M, Sullivan GJ, Bindoff LA, et al. Comparing the mitochondrial signatures in ESCs and iPSCs and their neural derivations. Cell Cycle. 2022;21(20):2206–21.

24. Mathapati S, Siller R, Impellizzeri AA, Lycke M, Vegheim K, Almaas R, et al. Small-Molecule-Directed Hepatocyte-Like Cell Differentiation of Human Pluripotent Stem Cells. Curr Protoc Stem Cell Biol. 2016;38:1g.6.1–g.6.18.

25. Siller R, Greenhough S, Naumovska E, Sullivan GJ. Small-molecule-driven hepatocyte differentiation of human pluripotent stem cells. Stem Cell Reports. 2015;4(5):939–52.

26. Siller R, Naumovska E, Mathapati S, Lycke M, Greenhough S, Sullivan GJ. Development of a rapid screen for the endodermal differentiation potential of human pluripotent stem cell lines. Scientific Reports. 2016;6(1):37178.

27. Strøm T, Martinussen T, Toft P. A protocol of no sedation for critically ill patients receiving mechanical ventilation: a randomised trial. Lancet. 2010;375(9713):475–80.

28. Liang KX, Chen A, Kristiansen CK, Bindoff LA. Flow Cytometric Analysis of Multiple Mitochondrial Parameters in Human Induced Pluripotent Stem Cells and Their Neural and Glial Derivatives. J Vis Exp. 2021 (177).

29. Wolf FA, Angerer P, Theis FJ. SCANPY: large-scale single-cell gene expression data analysis. Genome Biol. 2018;19(1):15.

30. Korsunsky I, Millard N, Fan J, Slowikowski K, Zhang F, Wei K, et al. Fast, sensitive and accurate integration of single-cell data with Harmony. Nat Methods. 2019;16(12):1289–96.

31. Asgrimsdottir ES, Arenas E. Midbrain Dopaminergic Neuron Development at the Single Cell Level: In vivo and in Stem Cells. Front Cell Dev Biol. 2020;8:463.

32. Zagare A, Barmpa K, Smajic S, Smits LM, Grzyb K, Grunewald A, et al. Midbrain organoids mimic early embryonic neurodevelopment and recapitulate LRRK2-p.Gly2019Ser-associated gene expression. Am J Hum Genet. 2022;109(2):311–27.

33. Xu C, Prete M, Webb S, Jardine L, Stewart B, Hoo R, et al. Automatic cell type harmonization and integration across Human Cell Atlas datasets. bioRxiv. 2023:2023.05.01.538994.

34. Braun E, Danan-Gotthold M, Borm LE, Vinsland E, Lee KW, Lönnerberg P, et al. Comprehensive cell atlas of the first-trimester developing human brain. bioRxiv. 2022:2022.10.24.513487.

35. Luo W, Brouwer C. Pathview: an R/Bioconductor package for pathway-based data integration and visualization. Bioinformatics. 2013;29(14):1830–1.

36. Mercurio S, Serra L, Nicolis SK. More than just Stem Cells: Functional Roles of the Transcription Factor Sox2 in Differentiated Glia and Neurons. Int J Mol Sci. 2019;20(18).

37. Osumi N, Shinohara H, Numayama-Tsuruta K, Maekawa M. Concise review: Pax6 transcription factor contributes to both embryonic and adult neurogenesis as a multifunctional regulator. Stem Cells. 2008;26(7):1663–72.

38. La Manno G, Gyllborg D, Codeluppi S, Nishimura K, Salto C, Zeisel A, et al. Molecular Diversity of Midbrain Development in Mouse, Human, and Stem Cells. Cell. 2016;167(2):566–80.e19.

39. Chen A, Yangzom T, Sullivan GJ, Liang KX. Mitochondrial Dysfunction and Neuronal Anomalies in <EM>POLG</EM> Mutant Midbrain Organoids. bioRxiv. 2023:2023.09.27.559684.

40. Liu G, Yu J, Ding J, Xie C, Sun L, Rudenko I, et al. Aldehyde dehydrogenase 1 defines and protects a nigrostriatal dopaminergic neuron subpopulation. J Clin Invest. 2014;124(7):3032–46.

41. Neuhoff H, Neu A, Liss B, Roeper J. I(h) channels contribute to the different functional properties of identified dopaminergic subpopulations in the midbrain. J Neurosci. 2002;22(4):1290–302.

42. Clemente-Suárez VJ, Redondo-Flórez L, Beltrán-Velasco AI, Ramos-Campo DJ, Belinchón-deMiguel P, Martinez-Guardado I, et al. Mitochondria and Brain Disease: A Comprehensive Review of Pathological Mechanisms and Therapeutic Opportunities. Biomedicines. 2023;11(9).

43. Anne Stetler R, Leak RK, Gao Y, Chen J. The dynamics of the mitochondrial organelle as a potential therapeutic target. J Cereb Blood Flow Metab. 2013;33(1):22–32.

